# Exploring the Hypothetical Role of *Bacteroides* Species in Depression Progression: Insights from Metagenomic Analysis

**DOI:** 10.1101/2024.11.20.624524

**Authors:** Zihua Li, Jingxi Sun, Jinlong Yang, Panpan Han, Liping Min, Yongkang Cheng, Yuanqiang Zou, Zhuhua Liu

**Affiliations:** School of Basic Medical Sciences, Ningxia Medical University, Yinchuan, Ningxia 750004, China; College of Life Sciences,University of Chinese Academy of Sciences, Beijing 100049, China; Mental Health Center, General Hospital of Ningxia Medical University, Yinchuan, Ningxia 750003, China; College of Forensic Science, Xi’an Jiaotong University, Xi’an, Shaanxi 710061, China; Laboratory of Genomics and Molecular Biomedicine, Department of Biology,University of Copenhagen, Copenhagen, Denmark

**Author notes:** Corresponding authors: Dr. Yuanqiang Zou, Dr. Zhuhua Liu. These authors contributed equally, Zihua Li, Jingxi Sun and Jinlong Yang contributed equally to this article.

**Keywords:** Depression, Metagenomics, Gut microbiome, *Bacteroides*, Random forest

## Abstract

Depression, a psychiatric disorder with significant morbidity and mortality, has a complex etiology. Recent advances in microbiome research have highlighted the potential role of gut microbiota in depression pathogenesis. This study utilized shotgun metagenomic sequencing to compare the fecal microbiota of 28 depression patients and 26 healthy individuals. Significant differences in gut microbiota composition were observed between the two groups. We generated 350 non-redundant high-quality metagenome-assembled Genomes (MAGs) by binning and conducted comparisons between the depression and control groups. Notably, we found that the MAGs enriched in people with depression mostly belonged to *Bacte*ndicating a close link between *Bacteroides* abundance and the development of depression, suggesting that *Bacteroides* might be a potential culprit for de*roides*, ipression. In the depression group, we found that the module of nitric oxide synthesis was remarkably enriched, and all *Bacteroides* MAGs were annotated to nitric oxide synthase, suggesting that increased levels of Bacteroides may contribute to elevated nitric oxide synthesis. Specifically, the mean relative abundance about the genomes of *Bacteroides xylanisolvens*, *Bacteroides caccae*, *Bacteroides fragilis*, *Bacteroides stercoris* and *Bacteroides ovatus* showed strong discriminatory power in distinguishing depressed patients from healthy individuals (AUC=0.834). This research shed light on the potential role of gut microbiota in depression and highlights specific metabolic pathways and microbial markers for further investigation.

**Improtance:** This research highlighted significant differences in the composition and function of gut microbiota between individuals with depression and healthy individuals, particularly the enrichment of *Bacteroides* MAGs in depression patients. The upregulation of the nitric oxide synthesis pathway associated with these MAGs belong to *Bacteroides* in the gut of depression patients had also been observed. The mean relative abundance of a specific set of *Bacteroides* MAGs had been identified, which could accurately classify individuals with depression from healthy individuals (AUC=0.834). Our results suggest the importance of exploring microbial markers as potential diagnostic and therapeutic targets in managing depression.

## Introduction

Depression, as a prevalent neuropsychiatric disease, is not only associated with high treatment costs, but also characterized by high incidence and mortality rates. It often undermines individuals’ psychological resilience and significantly increases the risk of suicide^1^. Globally, depression has a wide range of impacts, affecting approximately 322 million people, or 4.4% of the global population, with a prevalence rate exceeding the natural growth rate of the global population^2^. The number of people suffering from this disease is soaring globally, and its extensive influence has made it an important issue in the field of public health. According to relevant statistics, the lifetime prevalence of depression is as high as 15%, making it a major culprit endangering human health^3^.

The pathogenesis of depression is intricate and has yet to be fully elucidated, intricately intertwined with complex psychological and physiological reactions. Recently, Vinelli et al^4^ emphasized the pivotal role of the human gut microbiota in maintaining health, uncovering the extensive functionality of these tiny organisms, their similarity in function to host organs, and the depth of their interactions. Notably, the intimate connection between gut microbiota and the brain via the microbiota-gut-brain axis offers a novel perspective for understanding the pathophysiology of mental disorders, including depression^5^. Compared to healthy individuals, the gut microbiota composition in patients with depression is markedly distinct, characterized by an increased ratio of Bacteroidetes to Firmicutes, as evidenced by studies conducted by Zheng et al^6^ and Simpson et al7, underscoring the potential role of gut microbiota in depression pathology. By intricately modulating endocrine, immune, and neuroactive pathways, the microbiota profoundly influences the communication mechanisms between the gut and the brain. This process involves neurotransmitters such as γ-aminobutyric acid and catecholamines and metabolites like SCFAs and bile acids produced by microbes, which may trigger alterations in brain-derived neurotrophic factors^8^.

In the investigation of the relationship between depression and gut microbiota, traditional 16S rDNA sequencing has been widely applied. However, its limitation lies in identifying microbial taxa only up to the genus level, thus unable to precisely capture interspecies differences and functional or metabolic characteristics. To address this limitation, the introduction of metagenomics becomes paramount. Compared to 16S rDNA sequencing, metagenomics provides a robust tool for comprehensively deciphering the complex interactions between hosts and microbiota. Metagenomic assembly enables the reconstruction of draft genomes of bacteria, archaea, and viruses that are uncultivable in laboratory settings, facilitating strain-level gene and functional annotation, comparative genomic analysis, and evolutionary studies. This approach provides insights into the ecological adaptation mechanisms, nutritional interactions, metabolic functions, and other traits of these uncultivable strains. Moreover, it facilitates the investigation of key microbial species that play crucial roles in complex diseases, as well as the interaction mechanisms between pathogens and hosts, including their microevolutionary processes. At present gut microbiota in depression remain scarce. Building upon this backdrop, the present study employs metagenomic approaches to systematically dissect the gut microbiota structure of healthy individuals and patients with depression. It not only reveals significant differences in gut microbiota composition between the two groups but also identifies specific bacteria or microbial community structures associated with depression characteristics, thereby opening new avenues for a deeper understanding of the pathogenesis of depression.

In this study, we highlighted the significant differences in the composition and function of gut microbiota between patients with depression and healthy individuals, particularly the enrichment of *Bacteroides* MAGs in depressed patients. We also observed an upregulation of the nitric oxide synthesis pathway associated with these *Bacteroides* MAGs in the gut of depressed patients. The identification of a specific set of *Bacteroides* MAGs with an average relative abundance (AUC=0.834) accurately distinguishes individuals with depression from healthy individuals. Our findings underscore the importance of exploring microbial markers as potential diagnostic and therapeutic targets for treating depression.

## 2 Methods

### 2.1 Participant Selection

**Subject recruitment:** This study protocol was reviewed and approved by the Ethics Committee of the General Hospital of Ningxia Medical University (KYLL-2021-768). All participants or their legal guardians signed the informed consent form. The fecal samples were collected at the General Hospital of Ningxia Medical University. The study lasted from January 2022 to June 2023, during which time patients with depression at the Mental Health Center of Ningxia Medical University General Hospital were included in the study. The healthy control group was composed of senior students and graduate students from Ningxia Medical University. In this study, a total of 54 participants were recruited, of whom 25 were diagnosed with depression and 26 were healthy controls. The participants were aged between 18 and 49 years old, with a male-to-female ratio of 27:25. The patient’s depression was diagnosed according to the strict criteria of the Self-Rating Depression Scale (SDS), where a score of more than 53 indicates the presence of such a condition. The fluctuation range of SDS scores for patients with depression is 53.75 ∼ 112.5, and the average SDS score is 87.32. Collect fecal samples from all participants immediately and store them at −80°C.

**Inclusion criteria:** Case group: aged 18-60 years old. Outpatients who have first onset of the disease and have not taken any psychotropic drugs or have stopped taking them for two weeks or longer before enrollment. Meet or meet the diagnostic criteria for the corresponding diseases defined in the Diagnostic and Statistical Manual of Mental Disorders, Fifth Edition (DSM-V) and the International Statistical Classification of Diseases and Related Health Problems, Implementation (ICD-11).

**Normal group:** no record of any mental disorder or major negative life events, no history of alcohol or drug abuse.

**Exclusion Criteria:** Subjects who are under 18 years old or over 60 years old. Subjects with secondary mental disorders, such as those induced by certain medications or severe physical illnesses. Subjects with a history of alcoholism or drug abuse. Subjects with organic lesions or neurodegenerative diseases of the central nervous system, including stroke, Alzheimer’s disease, or Parkinson’s disease. Subjects with accompanying severe physical illnesses. Subjects with a history of head injury, brain parenchyma diseases, or brain tumors. Female subjects during menstruation, pregnancy, or lactation.

### 2.2 Stool Metagenomic Sequencing and Metagenomic Profile

We conducted shotgun metagenomic sequencing on fecal samples collected from 51 participants to obtain a comprehensive metagenomic profile. Initially, DNA extraction from the fecal samples was performed using the MagPure Stool DNA KF Kit55. Subsequently, shotgun sequencing was carried out on the DNBSEQ-T7 platform with 100bp paired-end reads. Following this, raw data from all samples underwent quality control by fastp(fastp --in11.fq.gz --in2 2.fq.gz --out1 1.fq.gz --out2 2.fq.gz --trim_poly_g --poly_g_min_len 10 --trim_poly_x --poly_x_min_len 10 --cut_front --cut_tail --cut_window_size 4 --cut_mean_quality 20 --qualified_quality_phred 15 --low_complexity_filter --complexity_threshold 30 --length_required 30, https://github.com/OpenGene/fastp) and removal of human-derived sequences to generate clean data by bowtie2(bowtie2 -q -p 16 --quiet -1 1.fq.gz -2 2.fq.gz -S sam -x GRCm39 --un-conc-gz filter.fq.gz, https://github.com/BenLangmead/bowtie2). Their average sequencing depth was 115606258.7.The clean data was then processed with kraken2 (kraken2 --use-names –db --threads 32 --report report.txt --gzip-compressed –output result.kraken fq.1.gz fq.2.gz, https://github.com/DerrickWood/kraken2) using default parameters to determine the microbial composition at the phylum, genus, and species levels.

### 2.3 Fecal Microbiome Functional Analysis

Functional analysis of fecal microbiomev was using gene set construction approach, with the following main steps: (1) Assembly of clean data into contigs using megahit (https://github.com/voutcn/megahit) with parameters −1 R1.fastq.gz −2 R2.fastq.gz -t 32 -o --out-prefix; (2) Gene prediction on assembled contigs for each sample using prodigal (https://github.com/althonos/pyrodigal) with parameters -m -p meta -i contigs. fa -f gff -o gff -d fna -s score.txt -a faa; (3) Merging predicted gene sequences from all samples and removing redundant sequences using cd-hit (https://github.com/weizhongli/cdhit) with parameters cd-hit-est -i faa -d 100 -c 0.95 -aS 0.8 -G 0 -M 0 -B 0 -T 10 -o fna, resulting in a non-redundant gene set containing 1,729,639 sequences; (4) Functional annotation of all sequences in the gene set after conversion to protein sequences using eggnog-mapper (https://github.com/eggnogdb/eggnog-mapper) with parameters emapper.py -i faa --output -m diamond --cpu 20 -d bact --data_dir eggnog5.0.2 --outfmt_shor. Linear discriminant analysis (https://github.com/SegataLab/lefse) was then used to identify all differentially expressed genes between depression patients and healthy individuals (LDA>2). Finally, omixer-rpm (java -jar omixer-rpm-1.1.jar -i result_matrix.tsv -d GBM.list -o -a 1 -c 0 -e 1, https://github.com/raeslab/omixer-rpm)was utilized to predict gut-brain modules for the differential genes in the two gut microbiomes.

### 2.4 Binning Metagenomic Data, Abundance Calculation of MAGs and Function Annotation of MAGs

We used binning tools metabat2 (default parameters, https://bitbucket.org/berkeleylab/metabat/src/master/), maxbin2 (http://downloads.jbei.org/data/microbial_communities/MaxBin/MaxBin.html), concoct(default parameters, https://github.com/BinPro/CONCOCT), After binning, we got a total of 12858 MAGs, Subsequently, we utilized CheckM2(default parameters, https://github.com/chklovski/CheckM2) to assess the quality of these MAGs, and based on the evaluation results, selected MAGs of high quality (completeness > 70%, contamination < 10%) for further analysis, which was 9,115 medium to high quality MAGs. Then 3759 MAGs were gained after removing redundant by drep software (-sa 0.99, https://github.com/maxpert/drep). Next, we clustered of all MAGs by drep(-sa 0.95).Through all steps, 350 high quality MAGs were selected as a aggregation of MAGs ultimately. Then referencing the GTDB-tk database, all MAGs underwent taxonomy profiling using the classify_wf module with default parameters in the GTDB-tk Software (https://github.com/Ecogenomics/GTDBTk). The gtdbtk infer and the website (https://itol.embl.de/) were utilized to construct an evolutionary tree of the MAGs. Finally, a database containing all MAGs was constructed using the kraken2-build module with default parameters to determine the abundance of all MAGs.

### 2.5 Statistical Analysis

The sequencing data underwent Wilcoxon test for group differences, while other data were analyzed using analysis of variance or chi-square test, with statistical significance set at p < 0.05. Random forest prediction was executed using the randomForest package in R (V 3.1.1). Additionally, α diversity and β diversity analyses were conducted using the vegan package in R (V 3.1.1).

## 3 Results

### 3.1 Characteristics of Study Population

At the General Hospital of Ningxia Medical University, we rigorously collected fecal samples from 3 patients with mild depression, as indicated by a Self-rating Depression Scale score exceeding 53, and from 25 patients with major depressive disorder, with scores exceeding 73. Simultaneously, we collected fecal samples from 26 healthy volunteers, and the detailed SDS scores of the collected depression patients could be found in Table 1. Following shotgun sequencing of these fecal samples, we obtained a total data volume of 304G, with the largest sample data size being 9.2G and the smallest size being 3.8G. The average data volume per sample was approximately 5.67±0.15G. After removing human-derived genic data from the metagenomic data, the data volume reduced to 293G, with the largest sample data size at 8G and the smallest at 3.7G. The average data volume per sample was approximately 5.46±0.14G.

**Table 1:**
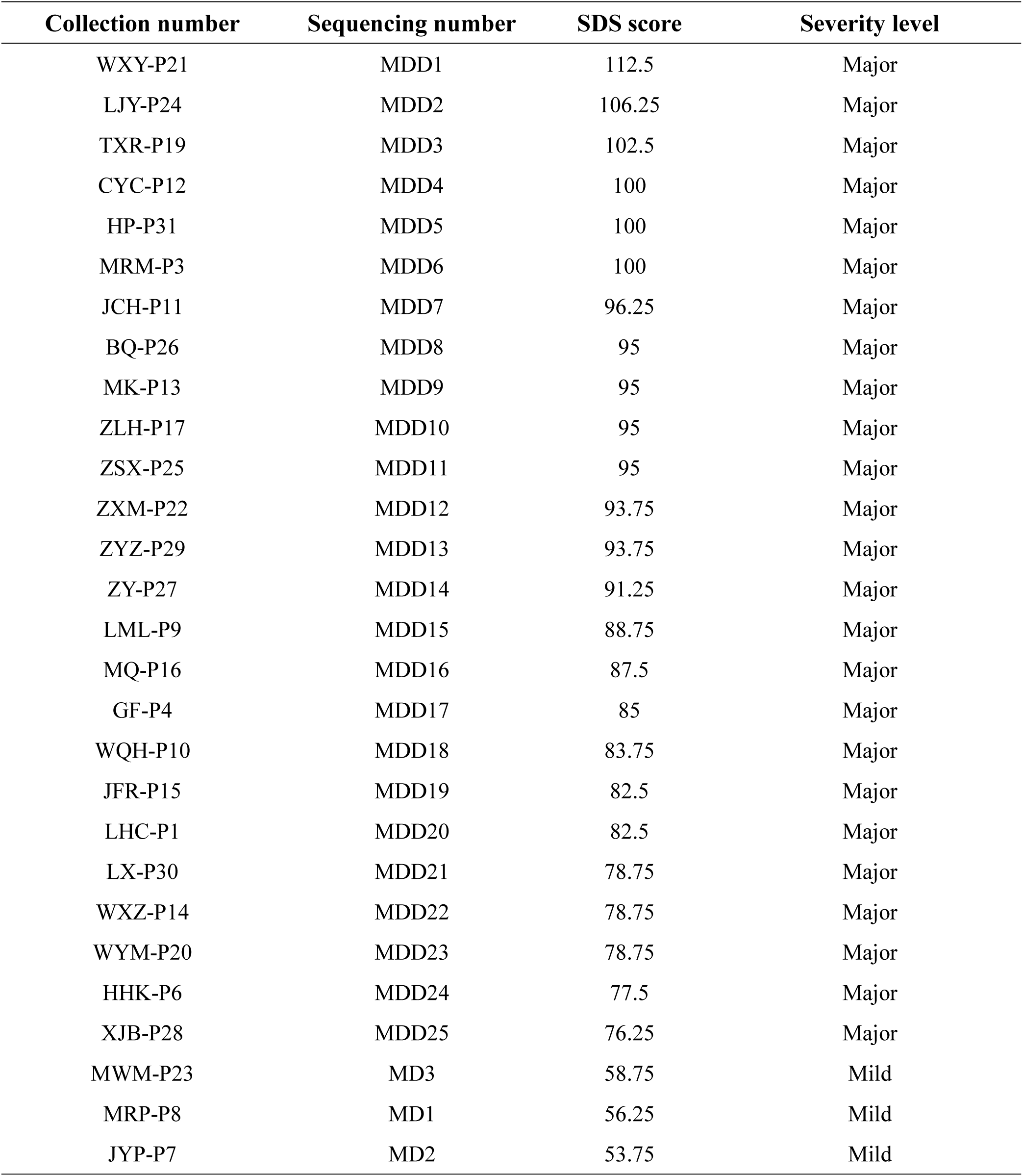
Detailed SDS scores of collected depression patients. This table provides detailed information on the Self-rating Depression Scale (SDS) scores of the collected depression patients. The table includes the sample ID, sequencing ID, severity level of depression, and the corresponding SDS score for each patient. The sequencing ID uniquely identifies each patient’s fecal sample. The severity level is categorized as either “Mild” or “Major” based on the SDS score.

| Collection number | Sequencing number | SDS score | Severity level |
| --- | --- | --- | --- |
| WXY-P21 | MDD1 | 112.5 | Major |
| LJY-P24 | MDD2 | 106.25 | Major |
| TXR-P19 | MDD3 | 102.5 | Major |
| CYC-P12 | MDD4 | 100 | Major |
| HP-P31 | MDD5 | 100 | Major |
| MRM-P3 | MDD6 | 100 | Major |
| JCH-P11 | MDD7 | 96.25 | Major |
| BQ-P26 | MDD8 | 95 | Major |
| MK-P13 | MDD9 | 95 | Major |
| ZLH-P17 | MDD10 | 95 | Major |
| ZSX-P25 | MDD11 | 95 | Major |
| ZXM-P22 | MDD12 | 93.75 | Major |
| ZYZ-P29 | MDD13 | 93.75 | Major |
| ZY-P27 | MDD14 | 91.25 | Major |
| LML-P9 | MDD15 | 88.75 | Major |
| MQ-P16 | MDD16 | 87.5 | Major |
| GF-P4 | MDD17 | 85 | Major |
| WQH-P10 | MDD18 | 83.75 | Major |
| JFR-P15 | MDD19 | 82.5 | Major |
| LHC-P1 | MDD20 | 82.5 | Major |
| LX-P30 | MDD21 | 78.75 | Major |
| WXZ-P14 | MDD22 | 78.75 | Major |
| WYM-P20 | MDD23 | 78.75 | Major |
| HHK-P6 | MDD24 | 77.5 | Major |
| XJB-P28 | MDD25 | 76.25 | Major |
| MWM-P23 | MD3 | 58.75 | Mild |
| MRP-P8 | MD1 | 56.25 | Mild |
| JYP-P7 | MD2 | 53.75 | Mild |

### 3.2 Gut Microbiota Disparities Between Depression Patients and Healthy Controls

The gut metagenomics data of 28 depression patients and 26 healthy individuals were analyzed to investigate potential differences in gut microbiota composition and structure between the two cohorts. We identified *Bacteroides, Phocaeicola, Faecalibacterium, Agathobacter, Segatella, Blautia* as the most prevalent taxa at the genus level in all individuals, however, there were big differences in the relative abundance of dominant bacterial genera among the samples (Fig1a). Subsequent Wilcoxon rank-sum tests at the phylum level demonstrated a notable rise in the relative abundance of *Bacteroidota* and a marked decline in *Bacillota* in the gut microbiota of depression patients in comparison to healthy controls (Fig1b and Fig1c). We used the Shannon and Simpson indices to assess the α-diversity of the gut microbiota at the genus level, and conducted principal coordinates analysis to reduce dimensionality and cluster the gut microbiota data, aiming to evaluate the structure of the gut microbiota at the genus level, the results showed significantly lower Shannon and Simpson indices in the gut microbiota of depression patients compared to healthy controls. Additionally, the structure of the gut microbiota at the genus level exhibited notable differences between depression patients and healthy individuals (Fig1d). These results emphasized clear compositional disparities in the gut microbiota between depression patients and healthy controls. An intriguing observation from our study was the absence of notable differences in the gut microbiota composition between individuals with mild depression and healthy controls (Sfig1). To mitigate potential interference in subsequent statistical analyses, samples from individuals with mild depression were excluded from the further study.

**Fig 1.**
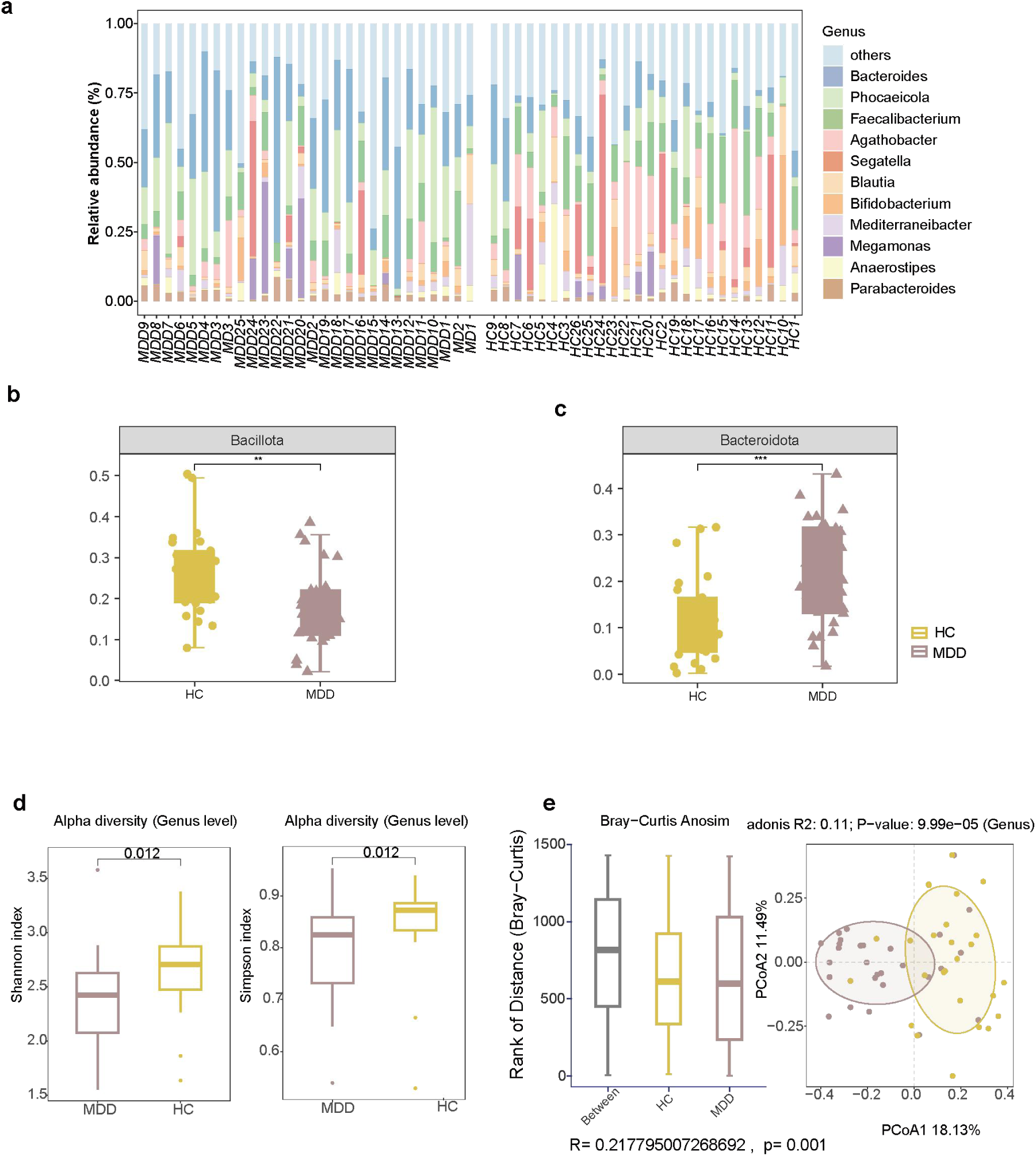
Gut microbiota composition analysis in depression people and healthy controls. (a) Stacked bar chart illustrating the taxonomic composition at the genus level for each sample. (b-c) Genera significantly enriched in the gut microbiota between depression patients and healthy controls (Wilcoxon test p < 0.05).(d)The difference in Shannon index and Simpson index at the genus level in the depression and control groups (Wilcoxon test p < 0.05).(e)The difference in β diversity by principal coordinate analysis at the genus level in the depression and control groups (PERMANOVA,p < 0.05). Yellow represents the healthy group, while purple represents the disease group

### 3.3 Differential Functional Profiles of Gut Microbiota Between Depression Patients and Healthy Individuals

In our effort to elucidate the functional difference of the gut microbiota in individuals with depression, we conducted a comprehensive exploration of microbial functions by building functional gene sets. Our methodology involved the genome assembly, gene prediction, and functional annotation of genes, culminating in the creation of a self-building and non-redundant functional gene sets. Subsequently, we quantified the expression levels of all functional genes in each sample and conducted principal co-ordinates analysis at the gene level to compare the two cohorts, unveiling notable differences in the functional profiles of the gut microbiota between individuals with depression and healthy controls (Fig2a). Following this, we identified the specific genes exhibiting differential expression between the two cohorts. Through chi-square tests, we identified 572 genes with remarkably elevated expression levels in the gut microbiota of individuals with depression, while 654 genes showed significantly increased expression in the gut microbiota of healthy individuals. To further understand the functional modules of these differentially expressed genes, we utilized the omixer-rmp tool to predict metabolic module profiles of the gut microbiota in both groups.

**Fig 2.**
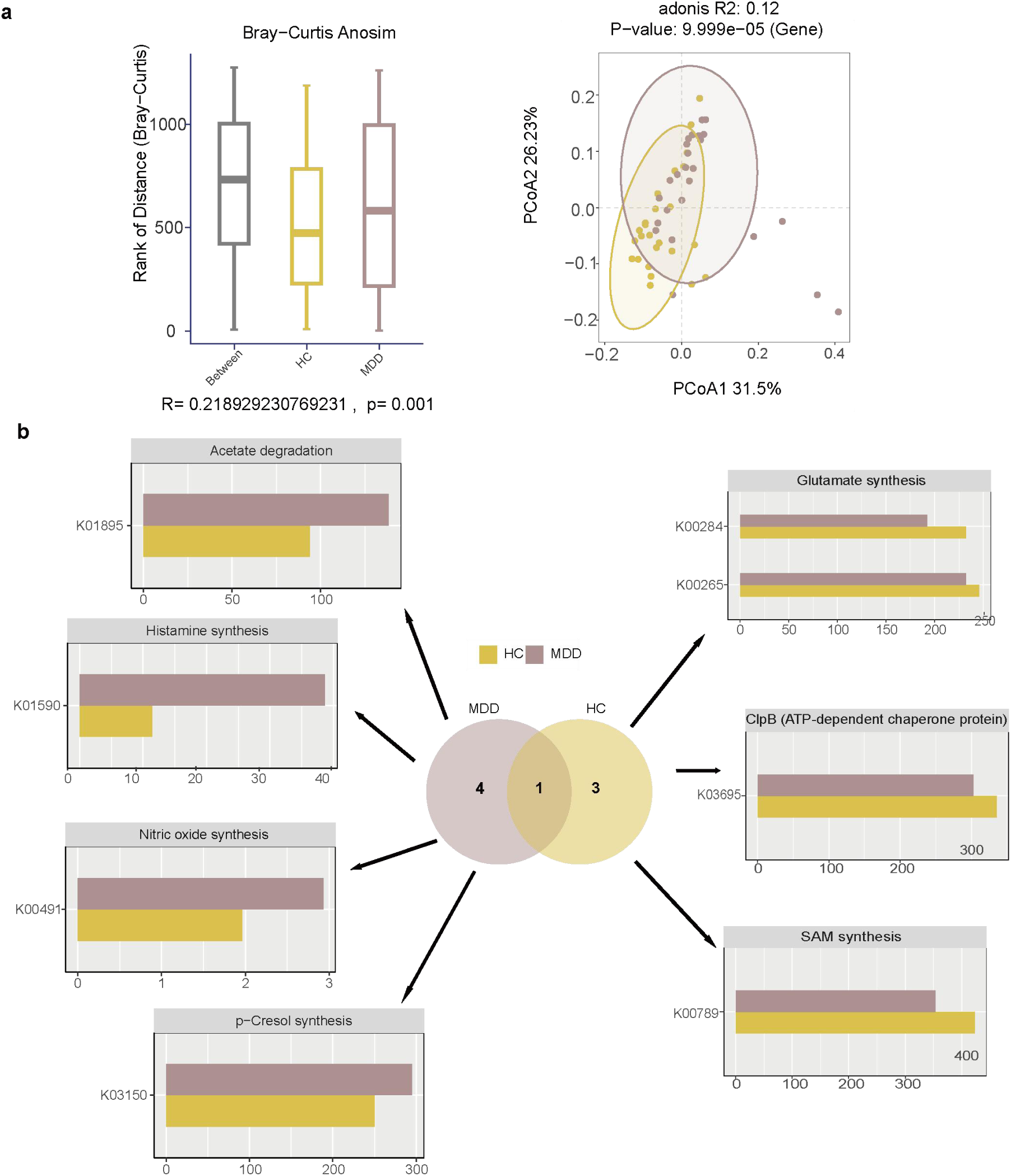
Gut microbiota functional analysis in depression patients and healthy people. (a)Left:the difference in Shannon index and Simpson index at the gene level in the depression and control groups (Wilcoxon test p < 0.05);Right:the difference in β diversity by principal coordinate analysis at the gene level in the depression and control groups (PERMANOVA,p < 0.05).(b)Metabolic module maps of the gut microbiota in both groups. The bar graph title indicates the gene’s gut-brain module and displays only genes with significant differences in the depression and control groups(Chi-square test p < 0.05). Yellow represents the healthy group, while purple represents the disease group.

As depicted in Fig1b, the distinct gene profiles of the gut microbiota in individuals with depression revealed specific metabolic modules including Acetate degradation, Histamine synthesis, Nitric oxide synthesis, and p-Cresol synthesis. Conversely, healthy individuals exhibited a different array of metabolic modules associated with Glutamate synthesis and ClpB (ATP-dependent chaperone protein) expression. Based on the aforementioned findings, it could be concluded that there ware significant difference in the gut microbiota between healthy individuals and patients with depression. The comparison of gene expression levels and the prediction of metabolic modules revealed distinct functional characteristics between the gut microbiota of the two groups.

### 3.4 Characterization of Enriched MAGs in Gut Microbiota of Healthy Individuals and Patients with Depression

In order to characterize the enriched bacteria in the gut microbiota of both healthy individuals and patients with depression, we conducted the assembly of all samples. Subsequently, the assembled contigs were binned by using three different tools (Concoct, MaxBin2, and MetaBAT2), leading to obtain of 9115 MAGs. Furthermore, after assessing the quality of the obtained MAGs by using CheckM2, we selected MAGs with a completeness of 70% or higher and contamination of 10% or lower. These MAGs, classified as intermediate to high quality, totaled 3759. Subsequently, we utilized the drep tool to acquire non-redundant MAGs, resulting in 350 MAGs. Then taxonomic annotation of these MAGs was conducted using the GTDB database, and the relative abundance of each MAG was calculated by the use of Kraken2. Phylogenetic evolutionary tree of these 350 MAGs was shown in Fig 3a, with detailed taxonomic annotation information provided in supplementary table 1. Finally, taxonomic annotation of these MAGs was conducted using the GTDB database, and the relative abundance of each MAG was calculated by the use of Kraken2. The results showed 31 MAGs enriching in the gut microbiota of healthy individuals and 19 MAGs enriching in the gut microbiota of patients with depression. The specific taxonomic information of these MAGs and their LDA values were presented in Fig3b. In the gut of healthy individuals, Enrichment was observed in multiple MAGs, including the MAGs annotated as *Agathobacter rectalis*, *Blautia_A wexlerae*, *Bifidobacterium adolescentis*, *Faecalibacterium prausnitzii*, *Gemmiger formicilis, Prevotella copri_A, Faecalibacillus intestinalis, Ligilactobacillus ruminis, Ruminococcus_E sp003526955, Mediterraneibacter faecis, Fimenecus sp000432435, Faecalibacterium sp, Hominisplanchenecus faecis, Holdemanella biformis, Dialister sp022722495, CAG-217 sp000436335, Coprococcus sp900066115, Faecousia sp900540635, UBA1417 sp003531055, CAG-103 sp000432375, Dialister sp900538805, Faecalibacterium prausnitzii_E, Eisenbergiella sp900066775, Blautia_A faecis, Romboutsia timonensis, Eubacterium_R sp000436835, Wujia chipingensis, Dorea formicigenerans, Blautia_A sp900066335, Faecalibacillus faecis* and *Faecalibacterium prausnitzii_I*. It has been confirmed that *Faecalibacterium prausnitzii* can prevent non-alcoholic fatty liver disease^9^, rheumatoid arthritis^10^, and atopic dermatitis^11^; oral administration of *Blautia wexlerae* can effectively slow down the progression of diabetes induced by a high-fat diet^12^, while *Bifidobacterium adolescentis* is a recognized producer^13^ of γ-aminobutyric acid which is a kind of neurotransmitter^14^ in the gut. Furthermore, many strains belong to the family *Lachnospiraceae*, several species of which have been shown to produce SCFAs, metabolites with beneficial effects derived from microbial metabolism. The MAGs including which were annotated as *Parasutterella excrementihominis, Bilophila wadsworthia, Bacteroides eggerthii, CAG-177 sp003514385, Enterocloster bolteae, Bacteroides, Bacteroides xylanisolvens, Bacteroides caccae, Bacteroides fragilis, Parabacteroides merdae, Phocaeicola massiliensis, Bacteroides fragilis_A, Parabacteroides distasonis, Phocaeicola coprocola, Bacteroides thetaiotaomicron, Bacteroides stercoris, Bacteroides ovatus, Bacteroides uniformis* and *Phocaeicola vulgatus* were enriched in the gut of patients with depression. Interestingly, in the gut of patients with depression, most of the enriched species taxonomized to the *Bacteroides* species, suggesting that changes in the abundance of species with *Bacteroides* might be associated with the development of depression. To elucidate the specific functions of each MAG enriched in the gut microbiota of healthy individuals and patients with depression, we conducted gene prediction and functional annotation on these MAGs sequentially. The final results were depicted in Fig4a. The gene K00491(nitric-oxide synthase), which responsible for carbon monoxide production, was found in all MAGs taxonomized into *Bacteroides.* Linear correlation analysis was conducted to examine the linear correlation between the abundance of these *Bacteroides* MAGs and the K00491 gene abundance, as shown in Fig 3b-c, B*acteroides stercoris* MAG*, Bacteroides thetaiotaomicron* MAG and *Bacteroides fragilis_A* MAG all exhibited significant positive linear correlations with the abundance of the K00491 gene. This suggested that the increased abundance of *Bacteroides* MAGs in the gut of patients with depression might lead to an elevation in nitric oxide synthesis. This finding aligned with the conclusion in 2.2, indicating a significantly enhanced capacity of gut microbiota in patients with depression to synthesize nitric oxide. The aforementioned research results underscored the dysregulation of the gut microbiota in individuals with depression compared to those who were healthy, and the enrichment of beneficial bacteria in healthy individuals could contribute to the maintenance of gut health and prevent depression, while the significant increase in *Bacteroides* species in patients with depression might be associated with the development of the disease.

**Fig 3.**
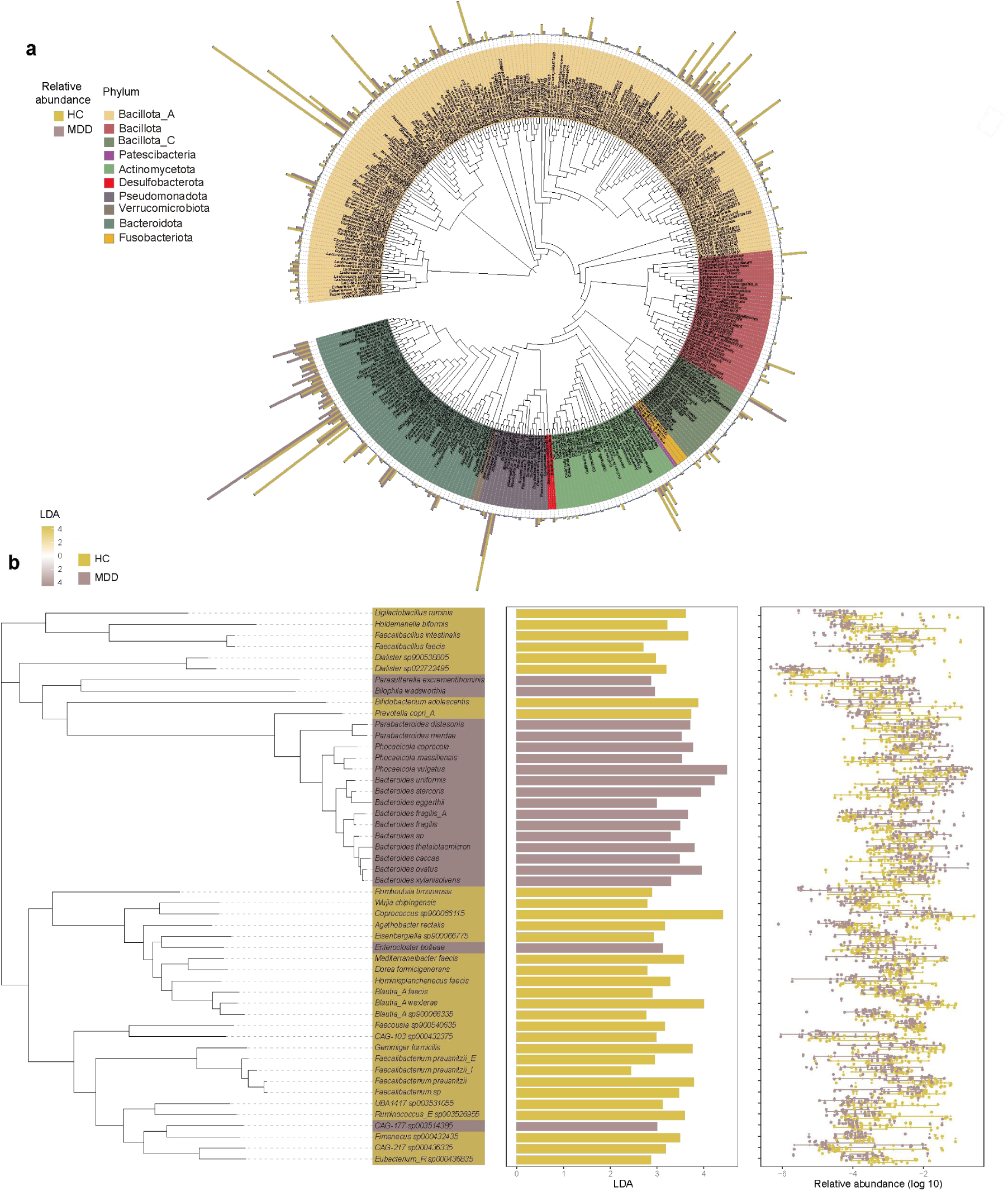
Comparative analysis of MAGs enriched in gut microbiota of depression patients and healthy ondividuals. (a) Evolutionary tree of MAGs, with different colors representing different phyla; (Left side of b) evolutionary tree of MAGs significantly enriched in the depression and control groups, with yellow indicating MAGs enriched in the gut of depression patients, and purple representing MAGs enriched in the gut of healthy individuals.(Right side of b) The bar graph represents the LDA values of two significantly different groups of MAGs, while the boxplot represents the relative abundance of MAGs.

**Fig 4.**
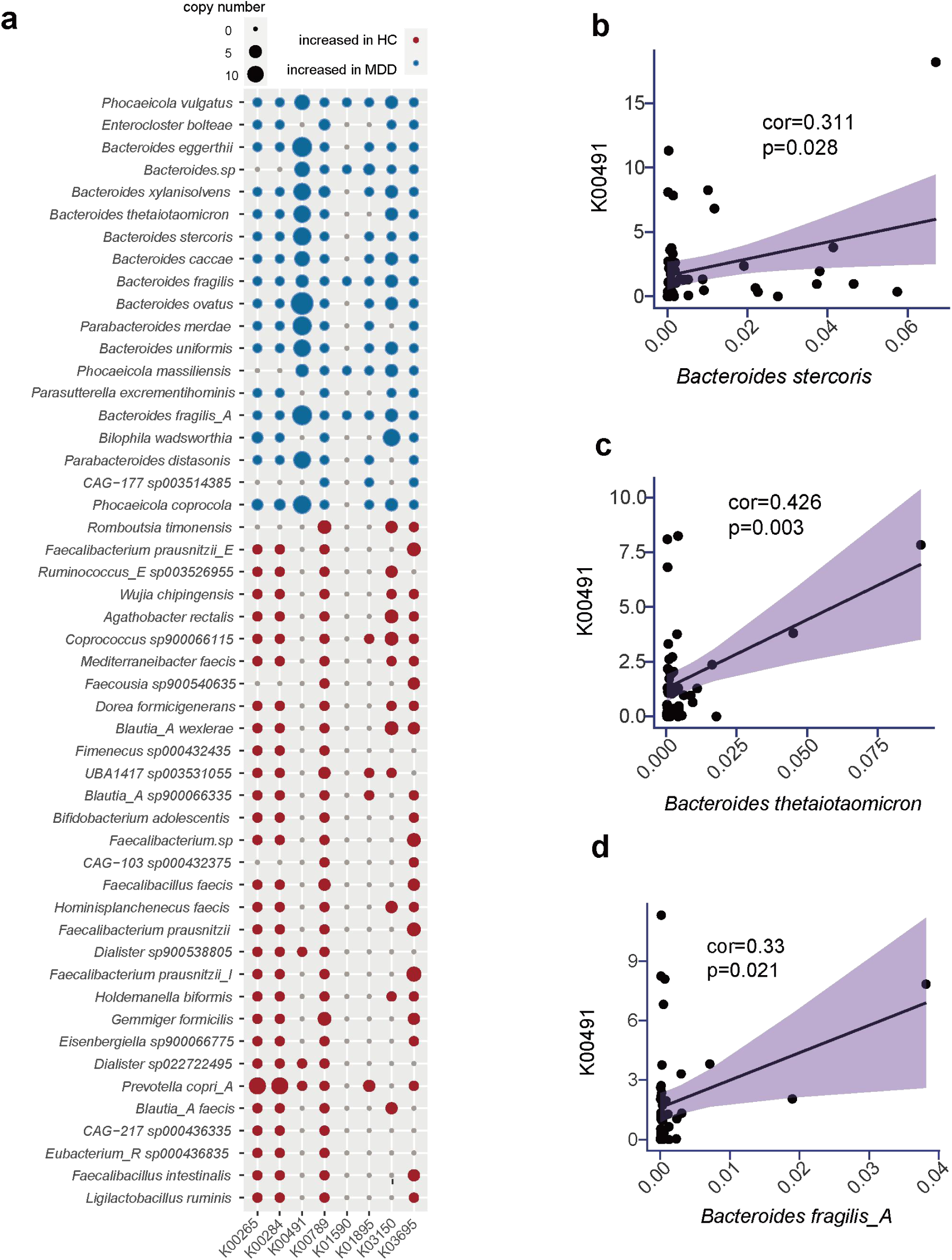
Gene prediction, functional annotation with enriched MAGs, and linear correlation analysis with gene K00491 abundance in *Bacteroides* species. (a) Gene prediction and functional annotation on MAGs significantly enriched in depression group and control group, with this figure only displaying genes consistent with those in Fig 2b; (b-d) Linear Correlation Results among Bacteroides stercoris, Bacteroides thetaiotaomicron and Bacteroides fragilis_A with Abundance of Gene K00491.

### 3.5 Analysis of the Importance of MAGs using Random Forest

Subsequently, we utilized a random forest model to investigate whether the enriched *Bacteroides* MAGs in the gut of patients with depression can distinguish between patients and healthy individuals. We selected the mean values of the combinations of *Bacteroides eggerthii* MAG*, Bacteroides sp* MAG, *Bacteroides xylanisolvens* MAG*, Bacteroides caccae* MAG*, Bacteroides fragilis* MAG*, Bacteroides fragilis_A* MAG*, Bacteroides thetaiotaomicron* MAG*, Bacteroides stercoris* MAG*, Bacteroides ovatus* MAG and *Bacteroides uniformis* MAG as predictive features, resulting in 1023 different feature combinations (refer to supplementary table 1). Then, we sequentially selected these features to determine the most effective ones for prediction (see the AUC of all features in supplementary table 2). The results, as depicted in Fig 5, showed that the mean values of *Bacteroides xylanisolvens* MAG*, Bacteroides caccae* MAG*, Bacteroides fragilis* MAG*, Bacteroides stercoris* MAG, and *Bacteroides ovatus* MAG (AUC=0.834) exhibited promising predictive performance. Similarly, the mean values of *Bacteroides caccae* MAG*, Bacteroides fragilis* MAG*, Bacteroides fragilis_A* MAG*, Bacteroides stercoris* MAG, and *Bacteroides uniformis* MAG (AUC=0.831) demonstrated notable predictive accuracy. Furthermore, the mean values of *Bacteroides eggerthii* MAG, *Bacteroides caccae* MAG*, Bacteroides fragilis* MAG*, Bacteroides fragilis_A* MAG*, Bacteroides stercoris* MAG and *Bacteroides uniformis* MAG (AUC=0.829) showed great predictive potential. The mean values of *Bacteroides caccae* MAG, *Bacteroides fragilis* MAG*, Bacteroides stercoris* MAG and *Bacteroides ovatus* MAG (AUC=0.82) also indicate predictive capability. Additionally, the mean values of *Bacteroides eggerthii* MAG*, Bacteroides caccae* MAG*, Bacteroides fragilis* MAG*, Bacteroides thetaiotaomicron* MAG*, Bacteroides stercoris* MAG and *Bacteroides uniformis* MAG (AUC=0.826) suggested predictive strength. Moreover, the mean values of *Bacteroides xylanisolvens* MAG*, Bacteroides fragilis* MAG*, Bacteroides stercoris* MAG and *Bacteroides ovatus* MAG (AUC=0.823) demonstrated predictive efficacy. Meanwhile, the mean values of *Bacteroides eggerthii* MAG*, Bacteroides fragilis* MAG*, Bacteroides stercoris* MAG and *Bacteroides ovatus* MAG(AUC=0.822) exhibit predictive potential. The mean values of *Bacteroides eggerthii* MAG*, Bacteroides caccae* MAG*, Bacteroides fragilis* MAG*, Bacteroides thetaiotaomicron* MAG and *Bacteroides stercoris* MAG (AUC=0.82) also showed notable predictive ability, which collectively demonstrated high discriminatory power. In conclusion, these findings suggested that variations in the abundance of *Bacteroides* MAGs could be utilized to predict the occurrence of depression.

**Fig 5.**
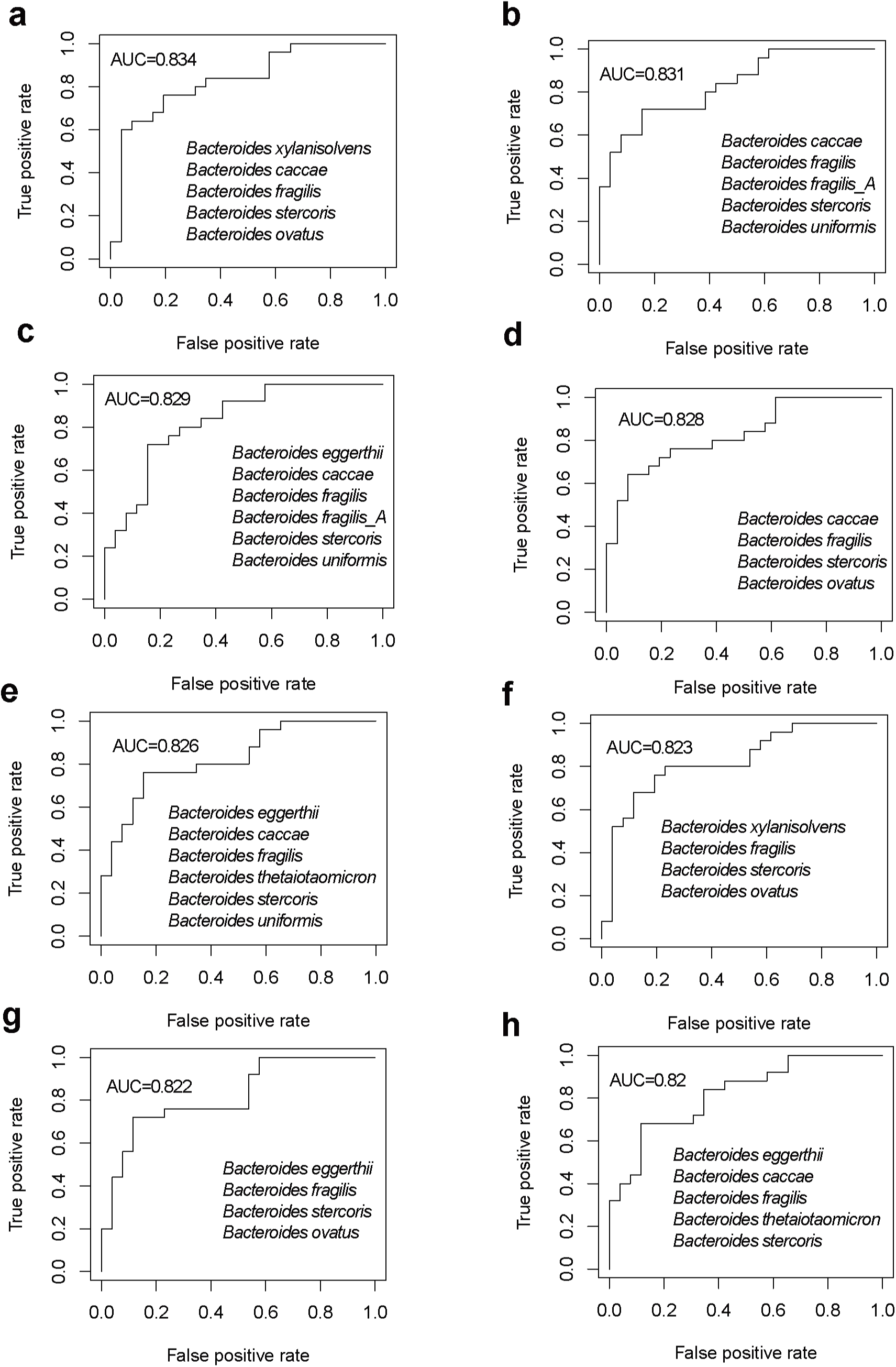
Random forest model analysis for predicting depression people based on enriched Bacteroides species abundance.

## 4 Discussion

This study employed shotgun metagenomic sequencing technology and metagenome-assembled genomes to systematically analyze the gut microbiota of patients with depression and healthy individuals, aiming to explore potential differences in gut microbiota composition and function between the two groups. The research findings revealed significant discrepancies in gut microbiota composition between patients with depression and normal individuals. Notably, a potential dysbiosis emerged in the gut microbiota of patients with depression, with changes in the abundance of *Bacteroides* MAGs, potentially closely associated with the progression of depression. This study not only contributes to a deeper understanding of the pathogenesis of depression but also has the potential to offer novel biomarkers and targets for the diagnosis and treatment of depression. By elucidating the relationship between gut microbiota and depression, we can further investigate the role of microbiota in neurological diseases, thereby providing a theoretical basis for developing interventions targeting the gut microbiota.

The pathogenesis of depression was complex, involving changes in both physiological and psychological aspects, and was mainly related to four factors: the hypothalamic-pituitary-adrenal (HPA) axis, the immune system, brain dysfunction, and the gut-brain axis^16,17^. The nearly 100 trillion bacteria in the human intestine affect the development of the immune system. The symbiotic microbiota can affect immune function, nutritional processing, and other physiological aspects of the host^18,19^. Recent studies have revealed the importance of intestinal microbiota in the function of the central nervous system, but the causes and consequences of the changes in intestinal microbiota caused by depression were still unclear^20^. Our research findings indicated that there were significant differences in gut microbiota composition between patients with depression and healthy controls. Notably, the relative abundance of *Bacteroides* significantly increases in the gut microbiota of depression patients, whereas the relative abundance of *Bacillota* significantly decreases. Furthermore, at the genus level, both the Shannon index and Simpson index of gut microbiota in depression patients were notably lower than those in healthy controls. Additionally, there existed a marked difference in β-diversity between the gut microbiota of depression patients and healthy individuals at the genus level. Previous similar reported on major depressive disorder have also found decreased Firmicutes and significantly increased levels of *Bacteroidota*, *Proteobacteria*, and *Actinobacteria*^21^. In a previous study, Yang et al. described a significant increase in *Bacteroides* in patients with major depressive disorder^22^. It can be seen that the decrease in the ratio of Firmicutes/Bacteroidetes is a common phenomenon in patients with depression.

Utilizing metagenomic approaches, we further assessed the unique gene expression profiles of the gut microbiota in patients with depression, revealing specific metabolic modules including acetate degradation, histamine synthesis, nitric oxide synthesis, and p-cresol synthesis. In contrast, the gut microbiota of healthy individuals exhibited distinct metabolic module arrays associated with glutamate synthesis and the expression of ClpB. This unique metabolic signature pattern observed in depression may be correlated with its pathogenesis. The altered gut microbiota structure in depression remodels the metabolic patterns of the microbiota, which in turn indirectly influences the metabolic or gene expression patterns of the host. The disturbed gut microbiota in depression exerts its influence on the gut nervous system through the secretion of metabolites, ultimately affecting brain function and contributing to the development of depression^23^. Previously, researchers reported that *Klebsiella* in the gut reduced serum estradiol levels through degradation by 3β-hydroxysteroid dehydrogenase, leading to depressive behavior in mice^24^. Similar vertical multi-cohort multi-omics joint studies have revealed that functional changes in gut microbiota (including various metabolic regulatory pathways such as proline metabolism) were closely related to the occurrence and development of depression^25^. It was evident that alterations in gut microbiota composition lead to shifts in microbial metabolic patterns, which were intimately linked to the onset of depression. This observation also suggested that modulating the metabolic patterns of the microbiome may emerge as a novel therapeutic direction for depression.

Our study employing shotgun metagenomic sequencing and metagenome-assembled genome techniques to explore the gut microbiome has yielded a more profound understanding in contrast to methodologies such as 16S rDNA gene amplicon sequencing or basic reads alignment for c analysis. This approach not only provides intricate genomic details of bacterial varieties but also enables the correlation of specific microbial genomes with gut-microbial-specific functions. For example, Hibberd, Matthew C et al.^26^ pinpointed *Prevotella copri* MAGs as a significant contributor to microbiome-directed complementary food (MDCF-2) induced alterations in metabolic pathways, particularly in the breakdown of polysaccharides found in MDCF-2. Another study delving into the functional aspects of the human gut microbiome revealed several uncultured species with the capacity to break down starch substrates^27^. In our research, by comparing the differences in gut microbiota MAGs between healthy individuals and those with depression, we found that one *Faecalibacterium prausnitzii* MAG was enriched in healthy individuals but significantly depleted in those with depression. *Faecalibacterium*, an anti-inflammatory species, was often downregulated in immune-mediated inflammatory diseases, and these associations may be mediated by the short-chain fatty acid butyrate^28^. It has also been reported that *Faecalibacterium* bacteria, which produce butyrate, were not observed in patients with depression^29^. *Faecalibacterium*^30^, *Ruminococcus*^31^, *Clostridia*^32^ and *Lachnospir*^33^ were the core microbiome that produces SCFAs in the intestine. Compared with normal individuals with depressive syndrome, there is a lack of microbiota that produce SCFAs, which are the products of certain intestinal probiotics and can serve as an energy source for colonic epithelial cells. These acids play crucial roles in the immune system, intestinal barrier, intestinal metabolism, and energy regulation. SCFAs can influence insulin resistance, ischemic stroke, hypercholesterolemia, and metabolic diseases^34,35^. In the gut of patients with depression, the majority of enriched MAGs belong to the genus *Bacteroides*, suggesting that variations in the abundance of *Bacteroides* species may be linked to the development of depression, particularly *Bacteroides xylanisolvens, Bacteroides caccae, Bacteroides fragilis, Bacteroides stercoris* and *Bacteroides ovatus*, the mean relative abundance of which exhibits the highest accuracy in distinguishing depressed patients from healthy individuals. Prior studies have reported an increase in the abundance of *Bacteroides* and a decrease in the abundance of Firmicutes and *Lactobacillus* in the gut microbiota of patients with depression. Stress-induced gut microbiota disturbances lead to reduced release of 5-hydroxytryptamine (serotonin) in the gut, impairing the functions of the prefrontal cortex and hippocampus, and resulting in decreased secretion of brain-derived neurotrophic factor (BDNF) and dopamine. These alterations contribute to emotional and cognitive impairments, ultimately triggering depression^36^. Our results indicated that the increased abundance of *Bacteroides* species may lead to enhanced nitric oxide synthesis. Previous studies have demonstrated that intestinal bacteria can secrete nitric oxide molecules, thereby altering the host’s own gene expression capabilities^35^.Relevant reports have indicated the dual nature of nitric oxide (NO) in organisms. Under physiological conditions, low concentrations of NO are produced via the L-Arg and constitutive NOS pathways, playing roles in signal transduction, maintaining vascular tone, and other physiological functions. However, under pathological conditions, the continuous production of large amounts of NO through the activation of inducible NOS exhibits cytotoxic effects. An appropriate amount of NO exerts neuroprotective effects, but an excess of it has neurotoxic effects^38,39^. Our research results indicated that the elevated level of *Bacteroidetes* MAGs in the gut of depressed patients leads to an increase in NO, which may cause direct damage to nerve cells or disrupt the homeostasis of the nervous system through indirect effects, thereby exacerbating the onset of depression. Further studies are required to reveal how the NO released by *Bacteroidetes* further aggravates the development of depression.

## 5 Conclusion

In summary, our study reveals significant alterations in the composition and function of gut microbiota in patients with depression compared to healthy individuals, with a notable enrichment of species belong to *Bacteroides* in individuals with depression. Furthermore, the involvement of *Bacteroides* MAGs in the nitric oxide synthesis pathway is significantly upregulated in the gut of individuals with depression. Using the average relative abundance of certain *Bacteroides* MAGs as a classification feature yielded precise classification results (AC=0.82∼0.834). Our findings suggest that *Bacteroides* species and the nitric oxide produced by gut microbiota may serve as effective targets for the treatment of depression.

## ACKNOWLEDGMENTS

This work has been supported by the Key R&D Program of Ningxia Hui Autonomous Region in China (2022BEG02033).

## AUTHOR CONTRIBUTIONS

Zihua Li, Conceptualization, Methodology, Writing – original draft | Jingxi Sun, Conceptualization, Methodology, Data curation, Software,Visualization, Writing–original draft | Jinlong Yang, Methodology, Writing – original draft | Panpan Han, Writing – original draft | Liping Min, Writing – original draft | Weiyi Mao, Writing – original draft | Yongkang Cheng, Writing–original draft | Yuanqiang Zou, Conceptualization, Project, Writing - review & editing | Zhuhua Liu, Conceptualization, Funding acquisition.

## DATA AVAILABILITY

The metagenomic sequencing data presented in the study are submitted in China National GeneBank (https://db.cngb.org/, project number: CNP0006487). This study does not provide code, please contact the corresponding author directly for access to the code. A STORMS checklist^40^ can be got at https://zenodo.org/records/14189203.

---

**Sfig1.**
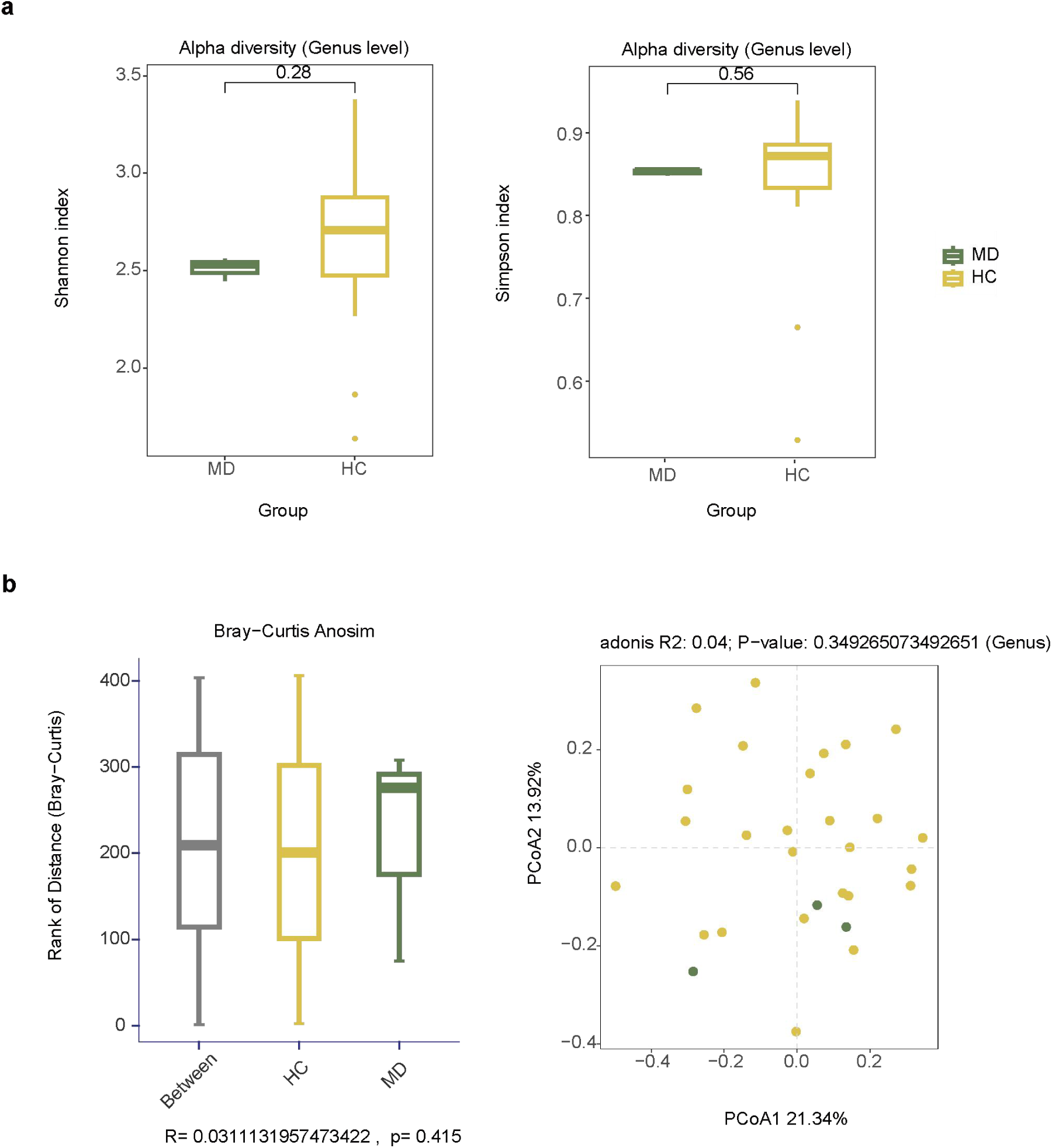
Gut microbiota composition analysis in mild depression people and healthy controls. (a)The difference in Shannon index and Simpson index at the genus level in the mild depression and control groups (Wilcoxon test p < 0.05).(b)The difference in β diversity by principal coordinate analysis at the genus level in the mild depression and control groups (PERMANOVA,p < 0.05). Yellow represents the healthy group, while green represents the mild depression group.

